# SPDEv3.0: A Multidisciplinary Integrated Data Analysis Platform

**DOI:** 10.1101/2025.03.30.646234

**Authors:** Dong Xu, Kangming Jin, Quanling Zhang, Xianjia Zhao, Yanchun Li, Tingkai Wu, Xiaobo Wang, Yuan Yuan, Zewei An, Zhi Deng, Wenguan Wu, Han Cheng

**Author notes:** Correspondence: Han Cheng,; Dong Xu. These authors contributed equally to this work (Dong Xu, Kangming Jin, Quanling Zhang, Xianjia Zhao, Yanchun Li).

## Abstract

Plant research encounters significant challenges in integrating multidisciplinary data and ensuring compatibility among diverse analytical methods. The limited automation in many existing tools further complicates data processing, thereby hindering efficiency and interdisciplinary collaboration. In response to these issues, we present SPDEv3.0, a multidisciplinary integrated data analysis platform specifically designed to tackle the complexities of data and the varied analytical requirements in plant sciences. Unlike traditional tools, SPDEv3.0 facilitates seamless cross-disciplinary data integration and automates analysis, featuring over 130 functions spanning plant breeding, molecular biology, genomics, and comparative genomics. The platform enables one-click operations for tasks ranging from data normalization to statistical analysis and visualization, significantly minimizing manual intervention and enhancing research efficiency. With its modular design and user-friendly interface, we believe that SPDEv3.0 will promote interdisciplinary collaboration and provides robust technical support for advancements in plant sciences.

## Introduction

Advances in plant science have ushered in an era of multidisciplinary research that integrates breeding science, molecular biology, genomics, and comparative genomics to address increasingly complex biological questions (Wang et al., 2024a; Yang et al., 2024). This shift has been driven by the need to uncover the genetic basis of traits, understand evolutionary relationships, and enhance crop performance in response to changing environmental conditions (Lyu et al., 2024). However, the integration of diverse data types and analytical methods presents significant challenges, particularly regarding data complexity, workflow automation, and accessibility for researchers with varying levels of technical expertise (Rose et al., 2021).

Despite the availability of numerous tools for individual tasks, many existing platforms remain inadequate for conducting multidisciplinary analyses within a unified environment. Data processing across various software platforms often encounters challenges such as data format conversion, complicating analysis and reducing its efficiency. Furthermore, certain disciplines still lack specialized software tailored for data analysis. For instance, breeding science heavily relies on genetic analysis; moreover, these tasks are still predominantly carried out using general-purpose statistical software such as SAS (Wang et al., 2024b). Since these tools are not specifically designed for breeding research, analyses often require multiple separate steps due to the lack of integrated and systematic functionalities. These unnecessary manual interventions reduce analytical efficiency and may introduce errors into the analysis process.

For example, in the case of the Two-Factor Randomized Complete Block Design (RCBD) analysis using SAS, researchers must individually perform the following steps: data preparation and import, analysis of variance (ANOVA) for two-factor randomized complete block design, post-hoc multiple comparison analysis, and residual analysis and normality testing. Since these steps are essential for completing RCBD analysis, integrating them into a single, comprehensive function could be a viable approach. Similarly, in genomics and comparative genomics analyses, certain multi-step processes have the potential to be integrated into a unified function. For example, in collinearity analysis, essential steps such as data extraction, sequence alignment, synteny identification, and visualization are all necessary. Therefore, consolidating these steps into a single streamlined function could significantly enhance analytical efficiency and eliminate potential data format conversion issues between different steps (Bernasconi, 2021).

To address the critical gap in analytical tools, we present SPDEv3.0, the latest iteration of the SPDE platform. Originally designed for molecular biology, this enhanced version now supports a broader range of disciplines, including breeding science, molecular biology, genomics, and comparative genomics. SPDEv3.0 integrates over 130 functions, with its most significant innovation being the reduction of unnecessary manual interventions across different analyses to further enhance automation. By streamlining workflows, it significantly improves analytical efficiency. For example, SPDEv3.0 can complete the collinearity analysis of *Arabidopsis thaliana* and *A. halleri* within one minute and design primers for 1,000 genes in just 33 seconds (Video S1 and S2). We believe that SPDEv3.0 will serve as a powerful tool for plant researchers, allowing them to focus more on data interpretation and analysis rather than expending effort on obtaining analytical results.

## Results

### Overview of SPDEv3.0

SPDE encompasses more than 130 functions, covering areas such as data analysis, file processing, statistical analysis, sequence preprocessing, and result visualization. These functions are systematically organized into seven modules, making them applicable across various disciplines, including molecular biology, genomics, comparative genomics, and plant breeding. An overview of the primary functions within each module is presented below (Figure 1).

**Figure 1.**
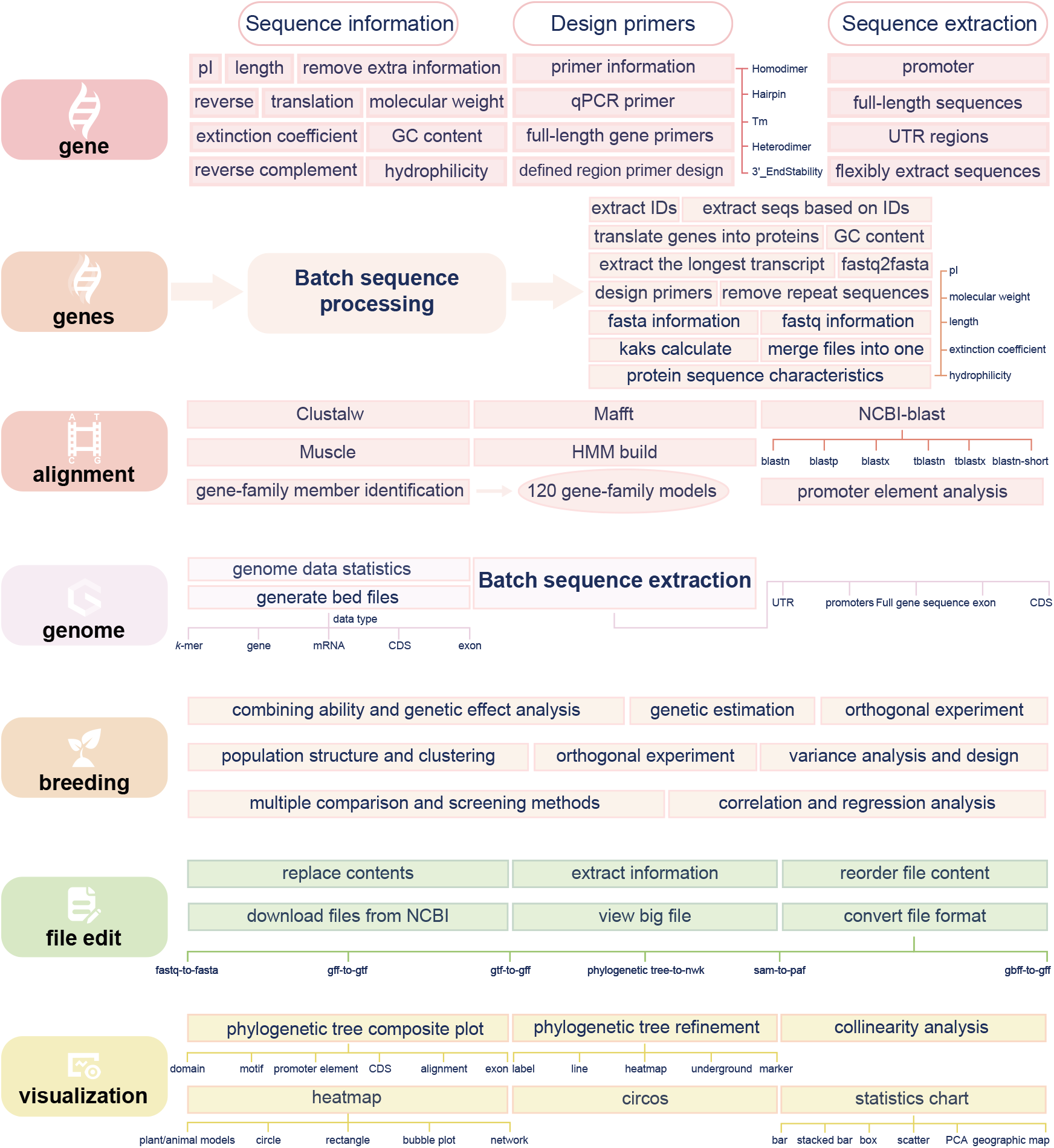
Overview of all SPDE functions.

Users can utilize the functions within the gene module to query basic sequence information, such as extinction coefficients and molecular weight, as well as to extract gene-related sequences, including promoters and untranslated regions, from the genome. In the area of primer design, users can easily design quantitative PCR primers and full-length gene primers, while specifying target regions for customized primer design. The functions within the multi-gene module are designed for batch processing tasks, encompassing bulk gene extraction, primer design, and gene translation. This module also facilitates the extraction of the longest transcript and provides statistical analysis of information from FASTA and FASTQ files. The alignment module includes multiple alignment tools, such as NCBI-BLAST, Diamond, ClustalW, MAFFT, and MUSCLE, all of which can be executed via user-friendly, click-based operations, thereby eliminating the need for command-line usage. Furthermore, the module allows users to generate corresponding Hidden Markov Models (HMMs) by inputting conserved domain sequences, which can be used to identify genes with conserved domains through HMMER’s hmmsearch function. Recognizing that searching for gene family members can be cumbersome, we have collected characteristic domains specific to each family and build relevant models that facilitate gene family analysis with a single click (for details, see the second section of “New Features and Key Improvements in SPDEv3.0”, Video S3).

Batch sequence extraction, including promoters, full-length genes, untranslated regions (UTRs) and coding sequences (CDS), is incorporated into the genome module. Additionally, BED files can be conveniently generated to display the distribution of elements, such as k-mers, genes, and mRNAs, across the genome. In the breeding module, we have incorporated various computational methods, including broad-sense heritability analysis, genetic correlation analysis, and orthogonal experimental design analysis to address data analysis challenges in plant breeding. After replacing the example datasets with their own, users can analyze the data with a simple click. In the file editing module, users can perform tasks such as content replacement, information extraction, data reorganization, format conversion, downloading files, and previewing large files. Data visualization has consistently been one of the most time-consuming processes; moreover, some tools require advanced coding and programming skills. To address these challenges, our visualization module integrates over 30 features for creating commonly used charts (details provided in the “Several Cases” section). It enables users to create charts through straightforward, click-based operations. Additionally, we have implemented extensive default parameters and automated processes to generate high-quality, publication-ready visualizations. For instance, we have developed 1,035 color schemes to automatically assign colors to plots generated by users. During the chart creation process, SPDE automatically performs data normalization, significance analysis, and automatic significance labeling (Video S4).

### New Features and Key Improvements in SPDEv3.0

#### 1. Automated computational methods for plant breeding

Although many gene functions and mechanisms have been confirmed and validated through experimental approaches such as molecular biology, and advancements in transgenic technologies have progressed rapidly (Lu et al., 2024), the development of new varieties still largely depends on traditional breeding techniques. In traditional breeding, breeders must determine whether traits can be reliably inherited through statistical analysis of the characteristics of new varieties. However, current breeding computations depend on statistical software such as SAS, and the calculation process remains relatively cumbersome. To enhance computational efficiency, we have designed functionalities for 40 breeding statistical methods (details in Supplementary Table S1) based on fundamental principles of breeding science (Acquaah, 2009; Chahal, G. and Gosal, 2002). We have included example data for each method, allowing users to easily replace it with their own and perform the corresponding calculations with a single click. Additionally, we have integrated plotting functions for some methods, enabling users to directly visualize the results.

#### 2. Streamlining processes for efficient genomic analysis

Genomic analysis is an inherently complex process often requiring multiple steps for completion. For instance, collinearity analysis between genomes involves several steps, including sequence alignment, gene location analysis, collinearity block identification, and visualization (https://github.com/tanghaibao/jcvi/wiki/MCscan-%28Python-version%29). Similarly, a basic analysis of gene families involves processes such as family member identification, sequence extraction, and domain analysis. The multitude of steps involved often reduces the efficiency of the analysis. Additionally, the need for multiple file format conversions, often caused by incompatible formats, increases the likelihood of errors and complicates the analysis process.

In SPDEv3.0, we have substantially optimized the genomic analysis workflow by automating critical steps that do not require user intervention, thereby enhancing both efficiency and reproducibility. This design enables users to perform complex analyses with a single click, significantly reducing manual effort and potential errors. To illustrate these advancements, we present examples of collinearity analysis and gene family analysis. In collinearity analysis, SPDEv3.0 automatically extracts coding sequences from genome sequences and GFF files provided by the user. The platform then translates the sequences and performs sequence alignments across multiple species. By systematically integrating annotation data from the GFF files, the software identifies collinearity blocks and generates comprehensive visualizations, effectively concluding the analysis. For example, SPDEv3.0 is designed to efficiently convert genomic files (*A. thaliana and A. halleri*) into collinearity visualization within one minute, demonstrating its capability for rapid and effective genome analysis (Video S1). Additionally, we assessed whether the efficiency of this function is correlated with genome size by analyzing genomes of varying sizes. The results indicate a strong correlation between runtime and genome size (correlation coefficient >0.96, Figure 2D); however, synteny analysis between large genomes (>2Gb) can still be completed within 2 minutes, demonstrating the high efficiency of this function.

**Figure 2.**
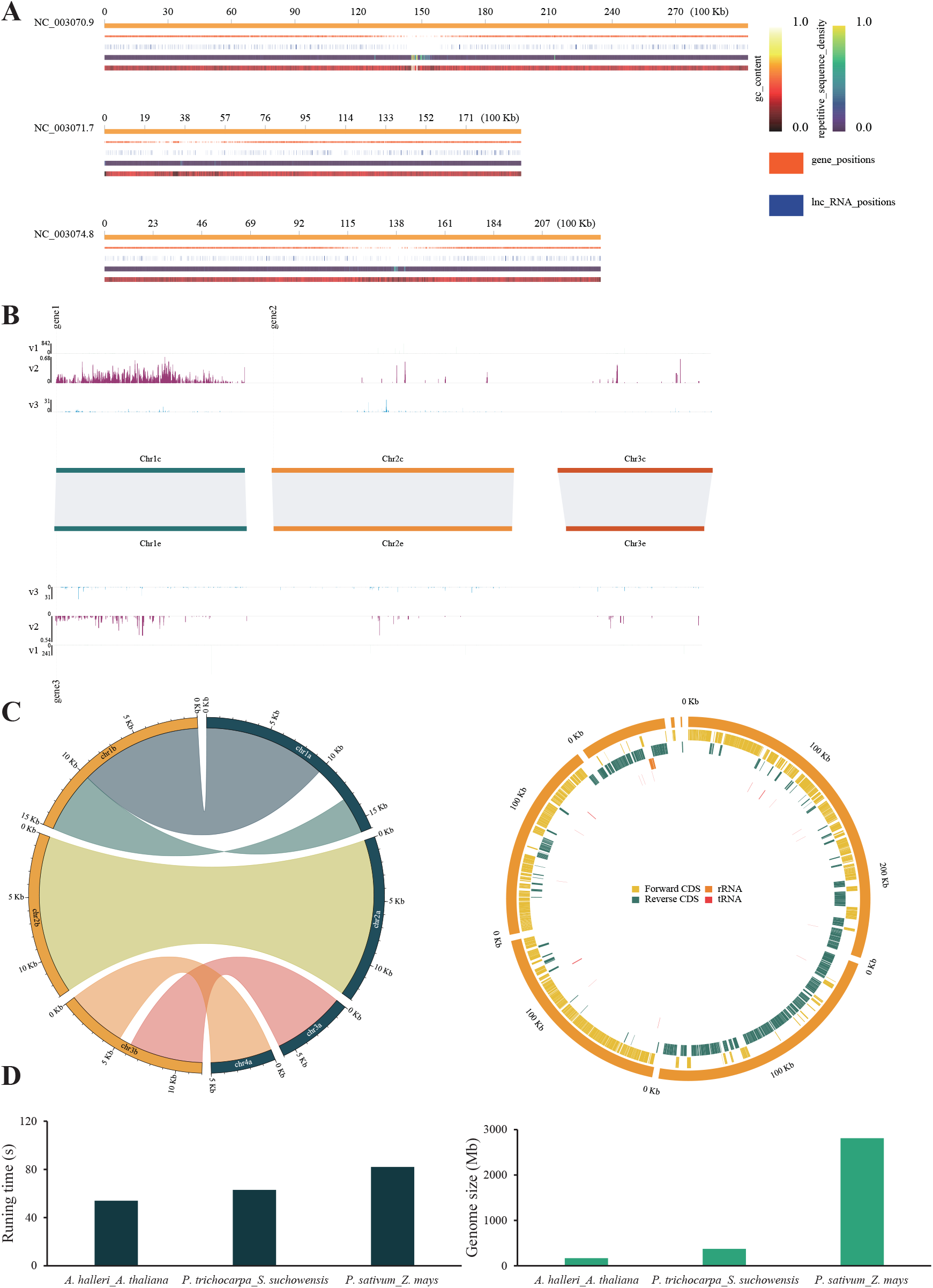
Genomic information visualization. **A**. Displaying genomic information. **B**. Comprehensive map of genomic synteny information and other data information. **C**. Circos plot. **D**. Statistical analysis of collinearity function efficiency. *P. trichocarpa, Populus trichocarpa; S. suchowensis, Salix suchowensis; P. sativum, Pisum sativum; Z. mays, Zea mays*

A similar automated approach has been employed for gene family analysis. In contrast to traditional methods requiring multiple tools to identify gene family members (Xu et al., 2022), SPDEv3.0 utilizes family-specific protein domain models (covering 126 gene families) to screen and preliminarily analyze family members. Using these models, the platform conducts a fully integrated workflow comprising family member identification, sequence extraction, domain recognition, and data visualization. For example, SPDEv3.0 can complete the entire process of family model selection, sequence extraction, domain analysis, and final domain model visualization for the maize auxin response factor (*ARF*) family within three minutes, demonstrating its efficiency in automated gene family analysis (Video S3).

#### 3. Batch processing enhances analytical efficiency

During gene family analysis and candidate gene screening, primer design for multiple genes is often required to determine their expression levels across different organs or under various treatment conditions. However, designing primers for multiple genes is a complex and time-consuming process. In SPDEv3.0, we have implemented a batch primer design function to efficiently streamline this process. Functional testing with 1,000 *A. thaliana* genes demonstrated that primer design for these genes can be completed within 33 seconds (Video S2).

Additionally, many analyses rely on sequences extracted from the genome. To facilitate this, SPDEv3.0 includes a batch extraction function that enables the retrieval of gene sequences, CDS, non-coding sequences, exon sequences, and promoter sequences across the entire genome. Using the Arabidopsis genome as an example, we found that our batch sequence extraction function can extract the CDS of all genes within four seconds (Video S5), significantly improving processing efficiency.

#### 4. Improvements in data analysis and operational convenience

Significance testing and data normalization are essential components of scientific research, particularly when substantial differences exist between datasets. Historically, these processes have been performed manually, requiring considerable time and effort. In the latest version of SPDE, we have automated these critical analytical steps, thereby enhancing both accuracy and efficiency. To further address the complexity of data analysis, SPDEv3.0 integrates a suite of commonly used analytical and visualization methods within its visualization module (see Figure 2 for examples). This comprehensive integration significantly streamlines workflows, facilitating more efficient completion of analytical tasks. SPDEv3.0 features a user-friendly interface that supports drag-and-drop operations and is available in both Chinese and English versions. Additionally, detailed help documentation and extensive example datasets are provided to assist users in navigating the platform effortlessly. These resources are designed to lower barriers to adoption and ensure that SPDEv3.0 is accessible to a broad user base.

### Several Cases

Compared to previous versions of SPDE (Xu et al., 2021), SPDEv3.0 introduces an expanded set of features for visualizing results. True datasets are utilized to demonstrate commonly used visualization types. As follows.

#### 1. Collinearity analysis and genomic information visualization

Collinearity analysis is essential in comparative genomics as it elucidates genome structure conservation, evolutionary trajectories, and syntenic relationships among species. This analysis facilitates gene mapping, genome assembly, and annotation, while providing insights into genetic relationships and adaptation. The genome information distribution module visualizes the positions of genomic elements on chromosomes, such as gene, lncRNA, GC content and repetitive sequence density, along with their quantitative distributions (Figure 2A). Meanwhile, this module also enables users to simultaneously visualize numerical data alongside genomic information, providing a more intuitive representation of differences across various genomic regions (Figure 2B). Furthermore, Circos plots are employed as an alternative approach to present more comprehensive and complex datasets (Figure 2C).

#### 2. Statistical graph

Scientific charting emphasizes statistical rigor, often incorporating significance testing (e.g., p-values, confidence intervals) to validate findings. These charts handle complex, multi-variable data and adhere to strict formatting standards set by academic journals to ensure clarity and reproducibility. We have automated critical processes such as significance analysis and data normalization to meet the specific requirements of scientific visualization in SPDEv3.0. This enables users to generate comprehensive data analysis and visualization results by simply inputting raw data. To accommodate the diverse visualization needs of researchers, the module includes a wide range of visualization methods such as volcano plots (Figure 3A), scatter plots (two and multiple variables, Figure 3F and B), PCA analyses (both 2D and 3D, Figure 3C and I), (Double) box plots (Figure 3E and D), (Stacked) bar charts (Figure 3G and H), violin plots (Figure 3J), bubble chart (Figure 3K), word clouds (Figure 3L) and sankey diagrams (Figure 3M). Significance analysis can be performed automatically for visualizations like bar charts, bubble charts, and word clouds. Additionally, visualizations such as PCA plots, box plots, and scatter plots support the simultaneous analysis and representation of multiple variables.

**Figure 3.**
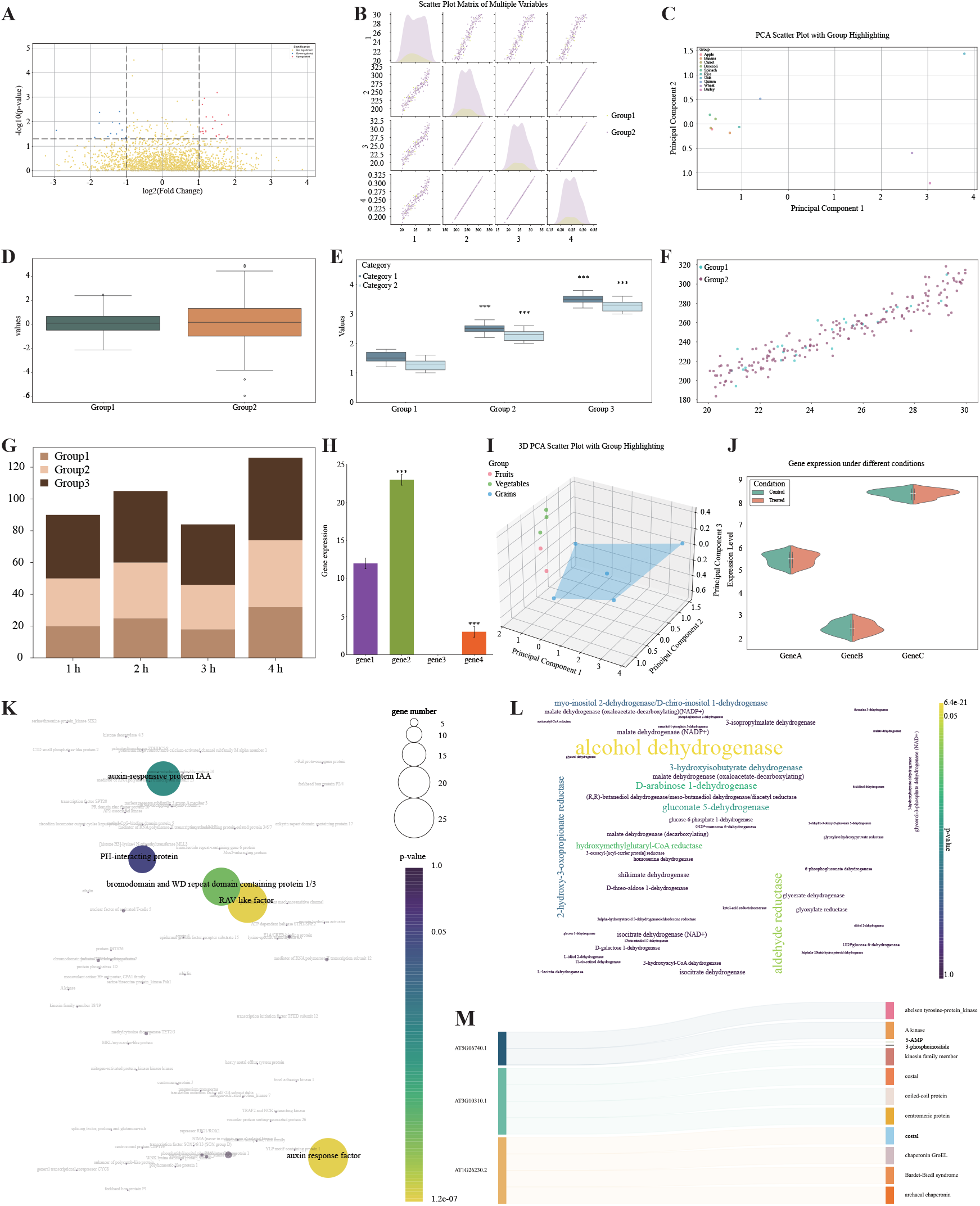
Statistical Chart. **A**. Volcano plot. **B**. Scatter plot matrix of multiple variables. **C**. 2D PCA plot. **D**. Box plot. **E**. Double box plot. **F**. Scatter plots of two variables. **G**. Stacked bar chart. **H**. bar chart. **I**. 3D PCA plot. **J**. Violin plot. **K**. bubble chart. **L**. Word clouds. **M**. Sankey diagrams.

#### 3. Heatmaps

Heatmaps are valuable tools for visualizing complex datasets, enabling the quick identification of gene-expression patterns, trends, and outliers through color-coded representations. They enhance interpretability and reveal relationships between variables that may be obscured in raw data. In SPDEv3.0, we have incorporated various heatmap formats, allowing users to select the most suitable one based on their data characteristics. Linear heatmaps are ideal for smaller datasets (Figure 4A), while circular heatmaps are better suited for larger ones (Figure 4D). Furthermore, SPDE allows combining circular heatmaps with phylogenetic trees (Figure 4B). Additionally, SPDE allows users to choose China or world maps (Figure 4E and F) and plant organs or cells (Figure 4C) to represent data more intuitively and effectively (Figure 4).

**Figure 4.**
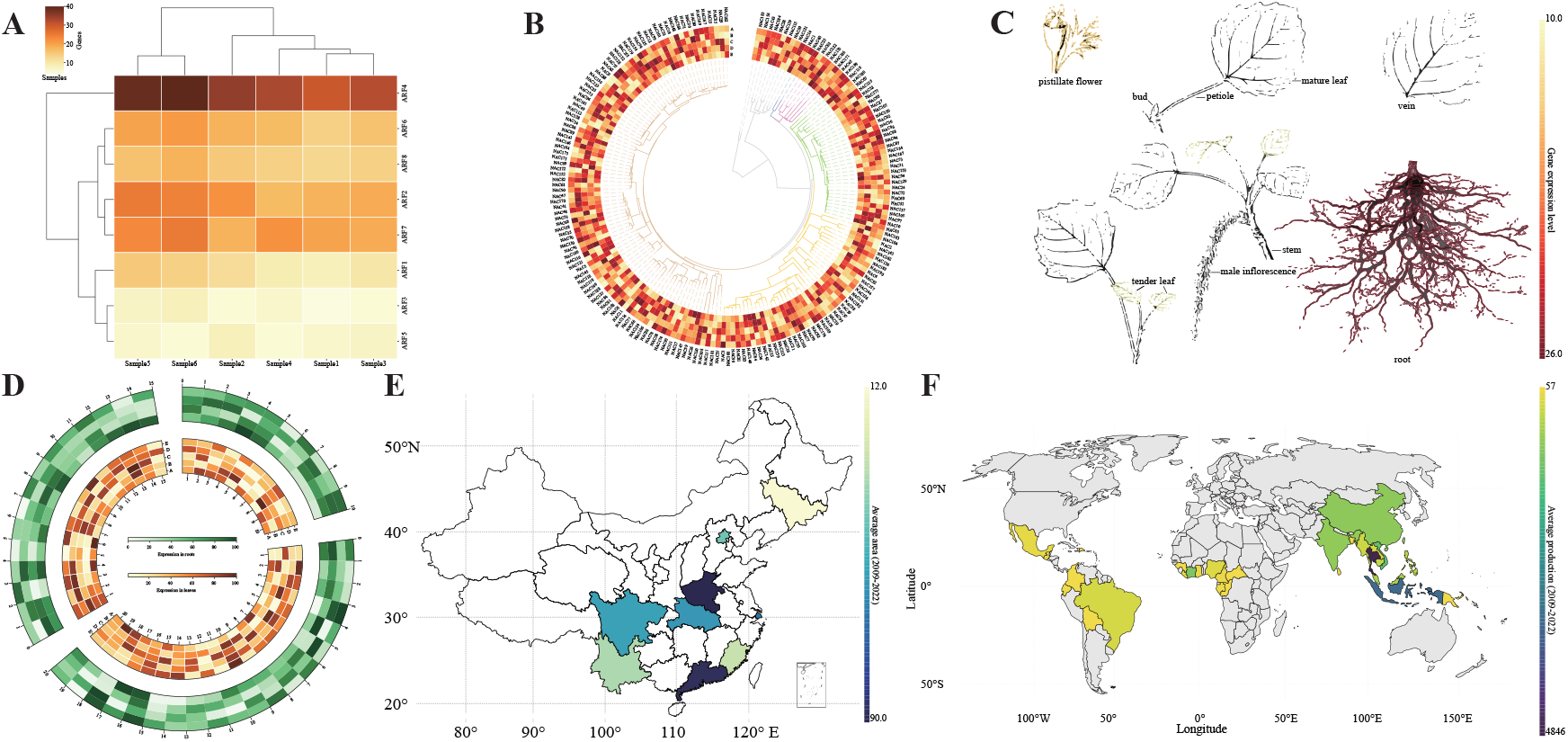
Visualization of data distribution (heatmap plot) **A**. Linear heatmap. **B**. Combining circular heatmap with phylogenetic tree. **C**. Plant organ plot. **D**. circular heatmap. **E**. China map. **F**. world map.

#### 4. Gene families

Gene family analysis is essential in plant research. It uncovers gene diversity, evolutionary mechanisms, and adaptation processes. Additionally, it identifies genes associated with traits such as disease resistance and yield, facilitating crop improvement and precision breeding. With the increasing availability of plant genomic data, this approach has become a fundamental tool for understanding plant biology (Noble et al., 2022; Pei et al., 2023; Tong et al., 2024). Building on this foundation, SPDEv3.0 enables automated visualization of protein domains through its gene family member identification feature (Figure 5A), illustrating conserved features. Conserved motifs within sequences can be analyzed using XML outputs from the MEME web tool (https://meme-suite.org/meme/tools/meme) (Figure 5B). Batch analysis and visualization of promoter elements are supported by inputting promoter sequences into SPDEv3.0 (Figure 5C). Additionally, SPDEv3.0 includes automated transcription factor binding site prediction and visualization, exemplified by *AP2* binding sites (Figure 5D). The platform also allows users to enhance the aesthetics of evolutionary trees for improved presentation (Figure 5E).

**Figure 5.**
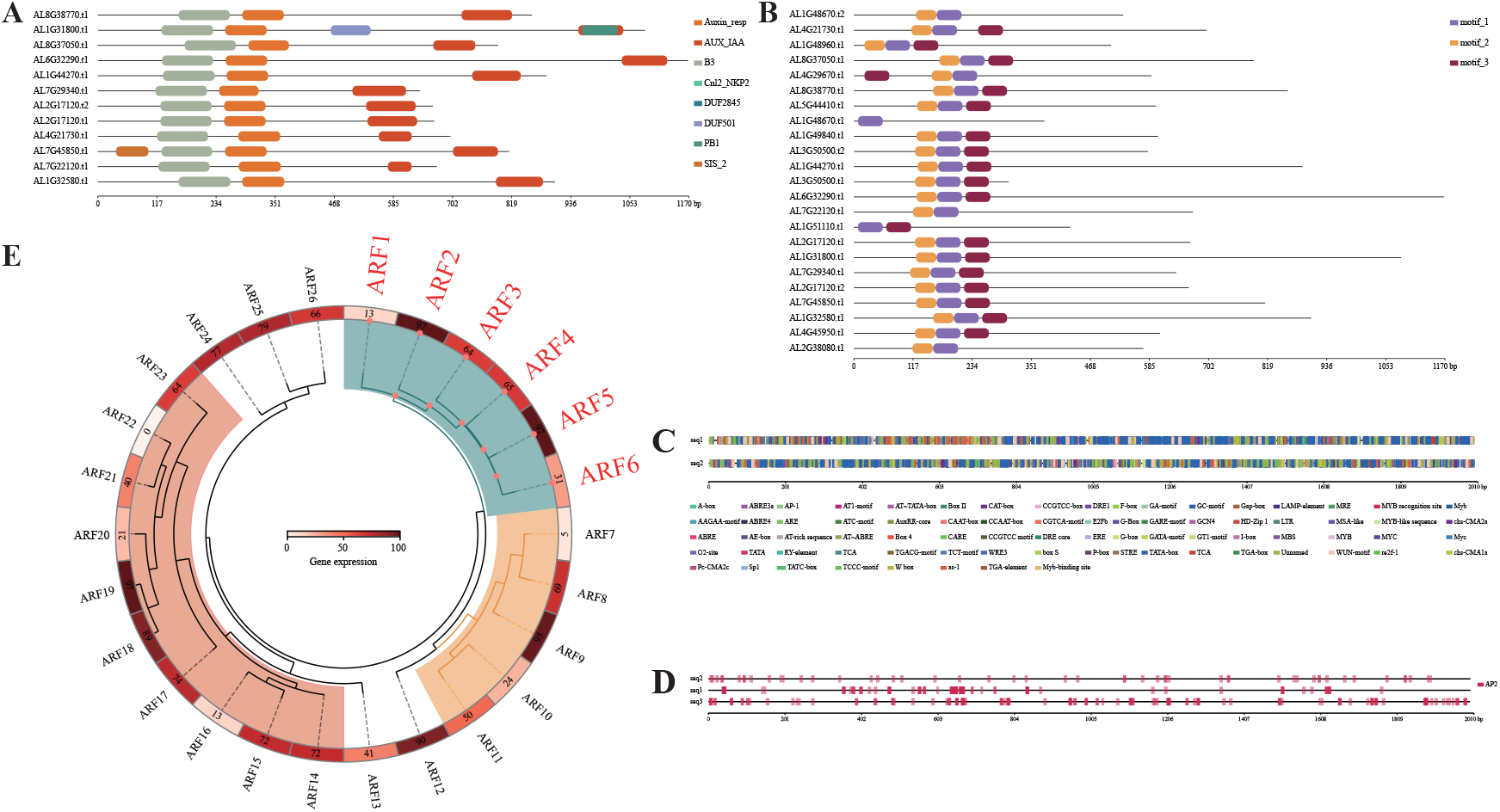
Visualization of gene family information. **A**. Protein domains. **B**. conserved motifs. **C**. Enhancement of phylogenetic tree visualization. **D**. Promoter elements. **E**. Interested element in promoter.

## Materials and Methods

### Data source

The genomic datasets used to evaluate SPDE’s functionality were derived exclusively from validated sources, primarily obtained from the NCBI RefSeq database. Similarly, Pfam annotations employed for protein domain and gene family identification were sourced from the InterPro-Pfam database (https://www.ebi.ac.uk/interpro/entry/pfam/#table). These datasets ensure the accuracy and reliability of the analyses conducted using SPDE.

### Construction of gene family identification database

Gene family members typically contain multiple protein domains, though not all of these domains are characteristic of the family. Following an extensive literature review on gene families, we selected 126 common gene families (Supplementary Table S2) and compiled and organized their characteristic domain Hidden Markov Models (HMMs). Finally, we defined the core protein domains for each family and compiled these domains into a dedicated database, enabling the automated identification of family members in SPDE.

### Calculate methods

SPDE includes default statistical methods for significance testing, such as the t-test, analysis of variance (ANOVA), and multivariate analysis of variance (MANOVA). For data normalization, the platform employs the z-score method as the default approach, with additional normalization techniques available for user selection.

### Python packages and software

We utilized various Python packages to enhance the accuracy and efficiency of data extraction and computation, and employed several Python frameworks to facilitate the interface design and aesthetic enhancement of SPDE. These packages and frameworks can be found in Supplementary Table S3.

### Transcription factor binding site prediction

We utilized the promoter extraction function of SPDEv3.0 to extract the complete promoter sequences of Arabidopsis, rice, maize, and poplar. These sequences were analyzed using PlantPAN 4.0 (Chow et al., 2024) to identify transcription factor binding sites (TFBSs). The identified binding sequences were then classified and summarized based on different transcription factors, forming a TFBSs sequence database. When utilizing these databases, SPDEv3.0 automatically selects the corresponding database file based on the user-specified transcription factor. It then performs sequence alignment using SeqMap (Jiang and Wong, 2008), followed by automated statistical analysis to identify binding sites. Finally, the binding sites are visualized for further interpretation.

### Data visualization

To address visualization needs, we implemented a Python-based program that converts input data into SVG vector graphics, ensuring scalability and optimal clarity. For color scheme selection, we curated 1,035 color palettes from contemporary publications, allowing users to switch effortlessly between modes with a single click. This flexibility enhances the interpretability of complex datasets.

### Plant and cell models

To ensure model accuracy and provide versatile visual representation, we collected 38 widely studied plant model images and two cell models (encompassing plant and animal cells) from specialized textbooks. These images were edited using Adobe Illustrator, organizing different organs or tissues into separate layers (examples shown in Figure S1). This layer-based approach enables users to overlay text or data onto specific organs or tissues, enhancing the precision and customization of visualizations.

### The logic and detailed algorithms of SPDEv3.0

Some common bioinformatics analyses require multiple steps, which do not necessitate researcher selection or manual intervention. From this perspective, these steps can be automated. Taking gene family analysis as an example, the fundamental steps include identifying family members, extracting sequences, conducting domain analysis, and visualizing results. However, current tools still require users to perform these tasks step-by-step, which hinders the efficiency of the analysis. We have optimized these steps for more streamlined analysis. In the following sections, we present the underlying logic and detailed algorithms behind some functions of SPDEv3.0.

### Automatic method for full-length gene primer design

In molecular experiments, it is often necessary to design full-length gene primers for cloning CDS of the target gene. However, the presence of homologous genes in the genome (e.g., members of the same gene family) poses challenges for primer design, as the terminal regions of these homologous genes may be identical or highly similar. Additionally, abnormal GC content in the terminal regions of genes can further complicate primer design. To address these challenges, we found that the UTRs (exhibiting higher sequence specificity) can used for designing the first-round primers. After amplifying the target sequence using the first-round primers, a second set of primers targeting the terminal regions of the gene can be used to effectively clone the full-length CDS. We have automated this primer design process. When users input the relevant genome files, SPDEv3.0 extracts the UTR and corresponding CDS regions from the genome and first designs the initial primer set based on these sequences. Subsequently, a second primer set is designed based on the terminal regions of the CDS (Video S6).

### Transcript analysis

Alternative splicing and other mechanisms can generate multiple transcripts from a single gene, a process that is a crucial aspect of plant research (Zhong et al., 2024). The transcript analysis feature in SPDEv3.0 initially extracts the genomic sequences of all genes, including the UTR regions, from the genome based on the GFF file. Once the user inputs the assembled transcriptome data, SPDE v3.0 aligns the gene sequences derived from the transcriptome to their corresponding genomic sequences. This process identifies the corresponding transcript for each gene based on alignment consistency and the proportion of the aligned region relative to the full-length gene. Furthermore, the analysis includes the identification of the longest transcript and the spatial genomic localization of the genes.

### Gene family analysis

The identification of family members, sequence extraction, domain detection, and domain visualization are fundamental operations in gene family analysis. In SPDEv3.0, when the user inputs a protein sequence, SPDE identifies the IDs of family members based on the selected family model (which can also be constructed and provided by the user) and outputs the positions of domains on the member sequences. The corresponding sequences are then extracted based on the member IDs, and visualizations are generated according to the domain positions. These operations can be finished with a single click (Video S3).

### Collinearity analysis

To conduct synteny analysis, the following steps are required: (1) obtain gene sequences, (2) perform sequence alignment, (3) identify syntenic blocks by integrating GFF annotations with alignment results, and (4) visualize the synteny analysis results. In SPDEv3.0, we have automated these processes, allowing users to perform collinearity analysis simply by providing the genome files (Video S1). Specifically, we first extract gene sequences from the genome and translate them into protein sequences. After translation, sequence alignment is conducted, followed by the construction of syntenic blocks. Finally, the syntenic blocks are visualized to facilitate interpretation.

## Discussion

Compared to previous software that required step-by-step operations (Wang et al., 2012), SPDEv3.0 introduces several key improvements to enhance the efficiency of data acquisition and analysis. In terms of file input, SPDEv3.0 only requires users to provide raw input files. For example, in collinearity analysis, the required protein sequence files are automatically extracted and translated by SPDE from genome sequence files and GFF annotations. This automation prevents errors caused by mismatched file contents. Regarding workflow automation, we have automated all steps that do not require user intervention. In data handling, SPDEv3.0 allows users to input entire folders, after which the software automatically iterates through the files and executes the selected operations. In data analysis, we have implemented automatic data normalization and significance testing. In visualization, SPDEv3.0 features automated layout adjustments and multiple color scheme options to reduce the time spent on manual color selection. Overall, one of the core design principles of SPDEv3.0 is to fully automate processes that do not require user decision-making. Functional tests using real datasets demonstrate the effectiveness of these systematic optimizations. This effectiveness is reflected in two aspects: first, the simplification of operational procedures, and second, a significant improvement in execution efficiency. We believe that these optimized designs will simplify the use of SPDE, making it more accessible and widely adopted by researchers, thereby contributing to the improvement of scientific data analysis efficiency. In the future, we plan to incorporate machine learning models and integrate broader, more diverse datasets to further enhance the efficiency and accuracy of data analysis.

## Data availability

The executables (Windows and MacOS), example data, tutorials are all available at https://github.com/simon19891216/SPDE.

## Author contributions

H.C. and D.X. designed the experiments. D.X. developed the SPDEv3.0 software. K.J. and Q.Z. were responsible for data collection and organization. X.Z. and Y.L. contributed to the development of part of the code. T.W., X.W., Y.Y., Z.A. and Z.D. evaluated the software’s functionality. X.D. wrote the manuscript. H.C. and D.X. revised the manuscript.

## References

Acquaah, G. (2009). Principles of plant genetics and breeding (John Wiley & Sons).

Bernasconi, A. (2021). Data quality-aware genomic data integration. Computer Methods and Programs in Biomedicine Update 1, 100009.

Chahal, G. and Gosal, S.S. (2002) Principles and Procedures of Plant Breeding: Biotechnological and Conventional Approaches. Narosa Publishing House.

Chow, C.-N., Yang, C.-W., Wu, N.-Y., Wang, H.-T., Tseng, K.-C., Chiu, Y.-H., Lee, T.-Y., and Chang, W.-C. (2024). PlantPAN 4.0: updated database for identifying conserved non-coding sequences and exploring dynamic transcriptional regulation in plant promoters. Nucleic Acids Research 52, D1569–D1578.

Hoang, N.V., Sogbohossou, E.O.D., Xiong, W., Simpson, C.J.C., Singh, P., Walden, N., van den Bergh, E., Becker, F.F.M., Li, Z., Zhu, X.-G., et al. (2023). The Gynandropsis gynandra genome provides insights into whole-genome duplications and the evolution of C4 photosynthesis in Cleomaceae. The Plant Cell 35, 1334–1359.

Jiang, H., and Wong, W.H. (2008). SeqMap: mapping massive amount of oligonucleotides to the genome. Bioinformatics 24, 2395–2396.

Lu, L., Delrot, S., Fan, P., Zhang, Z., Wu, D., Dong, F., García-Caparros, P., Li, S., Dai, Z., and Liang, Z. (2024). The transcription factors ERF105 and NAC72 regulate expression of a sugar transporter gene and hexose accumulation in grape. The Plant Cell 37 (1), koae326.

Lyu, X., Li, P., Jin, L., Yang, F., Pucker, B., Wang, C., Liu, L., Zhao, M., Shi, L., Zhang, Y., et al. (2024). Tracing the evolutionary and genetic footprints of atmospheric tillandsioids transition from land to air. Nature Communications 15, 9599.

Noble, J.A., Bielski, N.V., Liu, M.-C.J., DeFalco, T.A., Stegmann, M., Nelson, A.D.L., McNamara, K., Sullivan, B., Dinh, K.K., Khuu, N., et al. (2022). Evolutionary analysis of the LORELEI gene family in plants reveals regulatory subfunctionalization. Plant Physiology 190, 2539–2556.

Pei, T., Zhu, S., Liao, W., Fang, Y., Liu, J., Kong, Y., Yan, M., Cui, M., and Zhao, Q. (2023). Gap-free genome assembly and CYP450 gene family analysis reveal the biosynthesis of anthocyanins in Scutellaria baicalensis. Horticulture Research 10, uhad235.

Rose, Y., Duarte, J.M., Lowe, R., Segura, J., Bi, C., Bhikadiya, C., Chen, L., Rose, A.S., Bittrich, S., Burley, S.K., et al. (2021). RCSB Protein Data Bank: Architectural Advances Towards Integrated Searching and Efficient Access to Macromolecular Structure Data from the PDB Archive. Journal of molecular biology 433, 166704.

Tang, H., Krishnakumar, V., Zeng, X., Xu, Z., Taranto, A., Lomas, J.S., Zhang, Y., Huang, Y., Wang, Y., Yim, W.C., et al. (2024). JCVI: A versatile toolkit for comparative genomics analysis. iMeta 3, e211.

Tong, C., Jia, Y., Hu, H., Zeng, Z., Chapman, B., and Li, C. (2024). Pangenome and pantranscriptome as the new reference for gene family characterisation – a case study of basic helix-loop-helix (bHLH) genes in barley. Plant Communications, 101190.

Wang, J., Xu, D., Sang, Y., Sun, M., Liu, C., Niu, M., Li, Y., Liu, L., Han, X., and Li, J. (2024a). A telomere-to-telomere gap-free reference genome of Chionanthus retusus provides insights into the molecular mechanism underlying petal shape changes. Horticulture Research, uhae249.

Wang, Y., Wu, G., Huang, S., Ma, L., Fan, H., Zhang, R., and Zhou, Z. (2024b). Analysis of genetic parameters of growth and wood traits provides insight into the genetic improvement of Schima superba. Tree Genetics & Genomes 20, 8.

Xu, D., Lu, Z., Jin, K., Qiu, W., Qiao, G., Han, X., and Zhuo, R. (2021). SPDE: a multi-functional software for sequence processing and data extraction. Bioinformatics 37, 3686–3687.

Xu, D., Yang, C., Fan, H., Qiu, W., Huang, B., Zhuo, R., He, Z., Li, H., and Han, X. (2022). Genome-Wide Characterization, Evolutionary Analysis of ARF Gene Family, and the Role of SaARF4 in Cd Accumulation of Sedum alfredii Hance. 11, 1273.

Wang, Y., Tang, H., DeBarry, J.D., Tan, X., Li, J., Wang, X., Lee, T.-h., Jin, H., Marler, B., Guo, H., et al. (2012). MCScanX: a toolkit for detection and evolutionary analysis of gene synteny and collinearity. Nucleic Acids Research 40, e49–e49.

Yang, Z., He, F., Mai, Y., Fan, S., An, Y., Li, K., Wu, F., Tang, M., Yu, H., Liu, J.-X., et al. (2024). A near-complete assembly of the Houttuynia cordata genome provides insights into the regulatory mechanism of flavonoid biosynthesis in Yuxingcao. Plant Communications 5, 101075.

Zhong, Y., Luo, Y., Sun, J., Qin, X., Gan, P., Zhou, Z., Qian, Y., Zhao, R., Zhao, Z., Cai, W., et al. (2024). Pan-transcriptomic analysis reveals alternative splicing control of cold tolerance in rice. The Plant Cell 36, 2117–2139.

